# Connexin 40 deficiency alters the temporal profile of postictal oxygen dynamics following focal seizures

**DOI:** 10.64898/2026.09.21.750745

**Authors:** Donald G. Welsh, Jordan S. Farrell, G. Campbell Teskey, Anil Zechariah

## Abstract

Epilepsy is increasingly recognized as a disorder involving both neuronal and vascular dysfunction. While connexin signaling has been implicated in epileptogenesis, the contribution of vascular connexins to seizure associated cerebrovascular pathology remains poorly understood. Connexin40 (Cx40) is an endothelial gap junction protein that plays a crucial role in vascular communication and blood-flow regulation. Seizures induce dynamic changes in cerebral perfusion and oxygenation, including prolonged postictal hypoperfusion/hypoxia. To determine whether Cx40 influences postictal hypoxia following focal seizures, we examined seizure characteristics and postictal oxygen dynamics in Cx40 knockout (Cx40^−/−^) mice using an established focal hippocampal seizure model.

Electrically kindled seizures were elicited in wild-type and Cx40^−/−^ mice, and local hippocampal tissue oxygenation was continuously monitored before and after seizure induction. Seizure duration did not differ between genotypes, indicating comparable seizure severity. Interestingly, Cx40 deletion altered the temporal pattern of postictal oxygen recovery, producing greater early hypoxia and a delayed secondary rebound in pO_2_ despite similar peak oxygen levels and overall hypoxic burden.

These findings demonstrate that loss of Cx40 selectively alters the temporal profile of postictal oxygen dynamics without affecting seizure duration. Taken together, the results suggest that endothelial gap junctional communication contributes to postictal vascular recovery and identify Cx40 as a potential modulator of seizure associated neurovascular dysfunction.

## Introduction

Epilepsy affects more than 50 million people worldwide, and approximately one third of these individuals remain resistant to currently available treatments^1,2^. Recent research has provided evidence for the role of non-neuronal mechanisms including intercellular communication through gap junction channels and hemichannels formed by connexins, as key players in sustaining hypersynchronous activity, drive neuroinflammatory cascades, and promote epileptogenesis in experimental and clinical epilepsy^1,3,4^. Most work in this area has focused on astrocytic connexin 43 (Cx43) which influences extracellular ion and glutamate homeostasis, propagation of calcium waves, and modulates seizure generation and spread^2,3,5^. Pharmacological targeting of astrocytic Cx43 channels has been demonstrated to alter epileptiform activity and mimetic peptides directed against Cx43 is shown to reduce spontaneous seizures^3,5^.

Recent work has reframed epilepsy as both an electrical and vascular disorder, showing that brief focal seizures trigger severe, regionally confined postictal hypoperfusion and hypoxia due to arteriolar vasoconstriction leading to approximately 50% reduction in blood flow in rodents and a comparable perfusion deficit in epilepsy patients^6,7^. This effect was shown to be mediated by cyclooxygenase-2 (COX-2) activity and L-type calcium channels, as pharmacological inhibition of these pathways prevented the vascular insufficiency and rescued postictal motor and memory deficits^6^. Postictal hypoperfusion and hypoxia have also been proposed as a unifying mechanism for diverse seizure-related pathologies including postictal cognitive and learning deficits^8^, sudden unexpected death in epilepsy (SUDEP)^9^, and progressive functional and behavioral deficits^10^.

Because postictal hypoperfusion and hypoxia reflect disruptions in cerebrovascular homeostasis, mechanisms that regulate vascular signaling and coordinated blood-flow responses may influence the magnitude and duration of these abnormalities. In this context, vascular connexins are of particular interest due to their roles in coordinating intercellular signaling within the neurovascular unit and their established involvement in vascular pathology in the central nervous system. Despite these links, the extent to which specific vascular connexins shape the magnitude, duration, or regional distribution of postictal hypoperfusion/hypoxia remains largely unknown. Connexin40 (Cx40) is a major vascular connexin that is highly expressed in endothelial cells which in the cerebral circulation, forms an integral part of endothelial electrical coupling and conducted vasomotor responses^11^. We have previously shown that endothelial gap junctional signaling is crucial for maintaining neurovascular responses, and that dysfunctional endothelial gap junctional signaling resulting from a genetic deletion of Cx40 critically impairs blood flow homeostasis during cerebrovascular challenges such as stroke^12^.

Given that vascular connexins contribute to neurovascular coupling and cerebral blood flow homeostasis, alterations in Cx40 could plausibly modify seizure susceptibility and the severity of postictal vascular compromise. In this context, we investigated how loss of Cx40 influences postictal oxygen dynamics by applying the same focal seizure paradigm used by Farrell et. al.^6^, comparing seizure features and postictal outcomes in Cx40 knockout mice (Cx40^-^/^-^) with those in wild type controls. By integrating a well-characterized connexin knockout mouse with an established model for examining postictal vascular dysfunction, our investigation provides initial insight into how endothelial connexin 40 signaling may modulate both seizure expression and the severity of postictal hypoperfusion/hypoxia during epilepsy.

## Materials and Methods

### Experimental animals

All procedures were approved by the Animal Care Committee at the University of Calgary (AC20-0170) and were conducted in accordance with the Canadian Council on Animal Care guidelines and the Animal Research: Reporting In Vivo Experiments (ARRIVE) recommendations. All animals were male, 8–10 weeks of age, weighing 21–27 g, and were group-housed under a 12-hour light/dark cycle with ad libitum access to food and water. Cx40^−/−^ mice were obtained from Janis M. Burt (University of Arizona) and are available through The Jackson Laboratory (B6.129S4-Gja5tm1Paul/J; Stock No: 025697). The Cx40^−/−^ colony was established as described in Zechariah et al., 2020^12^. Given the minor, non-vascular effects of Cx40 and to minimize breeding and animal use in accordance with Animal Care and Use Committee regulations, C57BL/6J mice were used as controls^12^.

### Seizure stimulation and oxygen detection

Methods for inducing seizures and recording local brain oxygen in the hippocampus were performed as previously described with minor modifications (Farrell et al., 2016)^6^. C57BL/6J and Cx40^−/−^ knockout mice were chronically implanted with twisted bipolar electrodes (1.5 mm posterior, 0.8 mm lateral, and 1.5 mm ventral to bregma) and an oxygen-sensing optode (2.2 mm posterior, 2.5 mm lateral, and 1.8 mm ventral to bregma) in the dorsal hippocampus. Implants were secured to the skull with four stainless-steel screws (one serving as a ground) and dental cement. Twisted bipolar electrodes were constructed using Teflon-coated stainless-steel wire (CAT#791400, A-M Systems, Carlsborg, WA).

Mice were allowed a 1-week recovery period following surgery before stimulation. Seizures were elicited using kindling stimulation (1 second train of 60 Hz, 1 millisecond biphasic square pulses), which produces a brief afterdischarge (electrographic seizure). Mice received 5–10 stimulations above seizure threshold before full oxygen recordings were performed, as oxygen profiles are more variable during initial stimulation sessions. Once consistent oxygen profiles were obtained, oxygen was recorded before, during, and after seizures to capture the full duration of postictal hypoxia. The absolute partial pressure of oxygen (pO_2_) was measured using an implantable platinum-based fluorophore (Oxford Optronix, Abingdon, United Kingdom). Using this system, local brain oxygen can be recorded in awake, freely moving animals at a sampling rate of 1 Hz^6,13^. Following seizure stimulation, a transient dip in oxygenation occurs, followed by a rebound and a subsequent period of prolonged hypoxia throughout the postictal period. We compared this neurovascular response between groups to identify potential abnormalities and quantified differences using independent-samples t-tests.

## Results

Given the established role of Cx40 in vascular communication and cerebral blood flow regulation, we investigated whether genetic deletion of Cx40 alters postictal oxygen dynamics following induced focal seizures. Brief hippocampal seizures were elicited in Cx40^-^/^-^ and C57BL/6J control mice using the same kindling stimulation parameters. Seizure duration did not differ between groups (Figure 1A, B), which is important because the severity of postictal hypoxia is strongly correlated with electrographic seizure duration^6^. We measured local hippocampal oxygen levels in awake, freely moving mice and noted similar levels of baseline oxygen in each group. The first 10 minutes of recording were expanded for detailed examination of baseline, ictal, and immediate postictal pO_2_ (Figure 1D–G).

**Figure 1.**
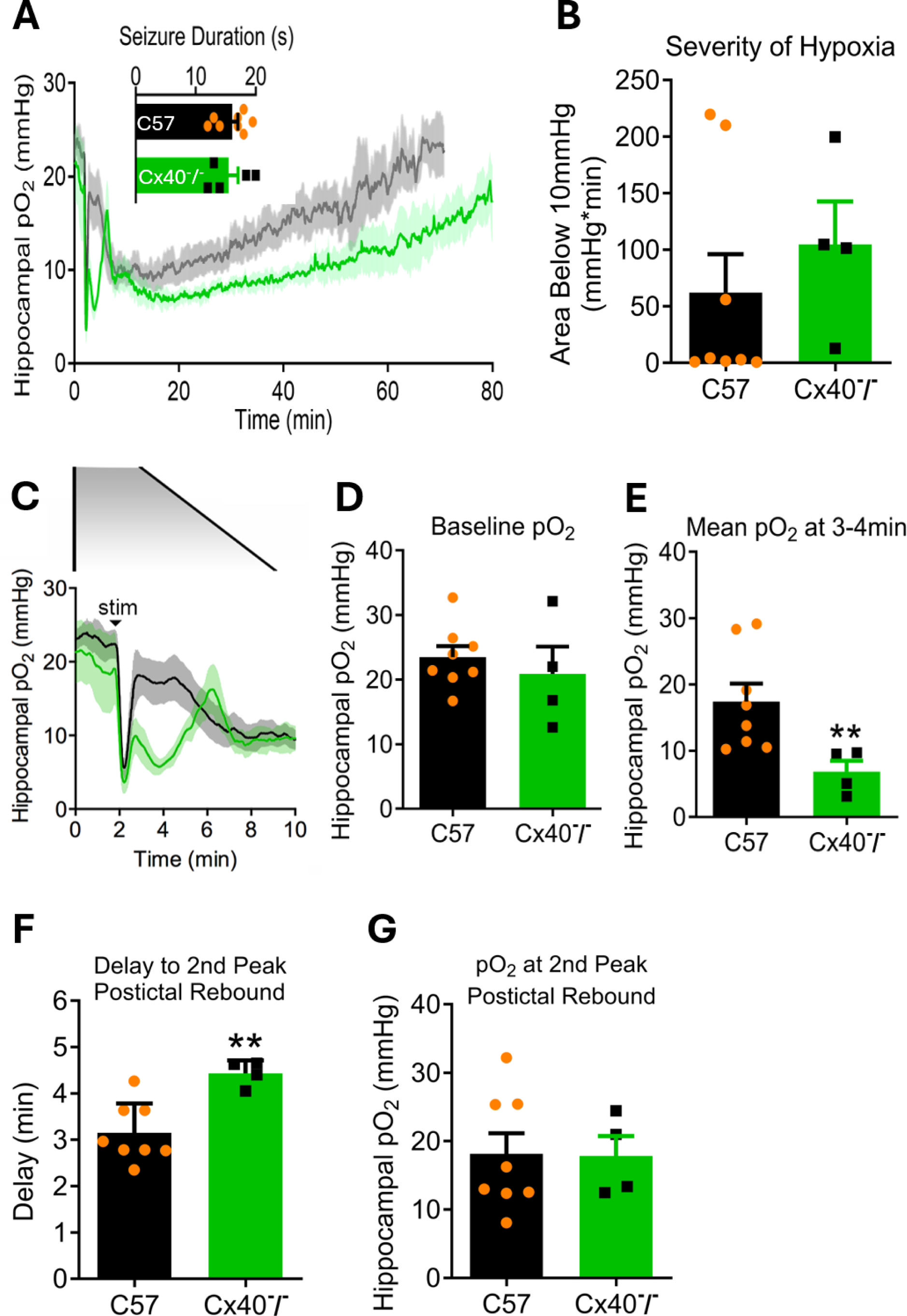
Cx40 knockout mice exhibit an altered neurovascular oxygen response following seizures. (A) Mean tissue oxygen tension (*pO*_2_) profiles recorded following kindled hippocampal seizures in C57BL/6J mice (black; *n* = 8) and Cx40 knockout mice (green; *n* = 4 – 5). Seizure duration did not differ between groups. The greatest divergence in *pO*_2_occurred during the immediate postictal period, which is shown at higher resolution in (C). (B) The overall severity of hypoxia, quantified as the area under the *pO*_2_curve below 10 mmHg, did not differ between groups. (C) Expanded traces showing the first 10 min of recording, including baseline, ictal, and immediate postictal *pO*_2_responses. Quantitative comparisons of these intervals are presented in (D–G). (D) Mean baseline *pO*_2_, measured over the 0–100 s preictal period, did not differ between groups. (E) Cx40 knockout mice exhibited significantly lower *pO*_2_than C57BL/6J mice during the 3–4 min postictal interval (\*\**p* = 0.0020, unpaired *t*-test). (F) Both groups exhibited a characteristic immediate postictal *pO*_2_response consisting of two distinct rebounds. The latency to the second postictal peak was significantly prolonged in Cx40 knockout mice compared with C57BL/6J mice (\*\**p* = 0.0061, unpaired *t*-test). (G) *pO*_2_at the second postictal peak did not differ between groups. Data are presented as mean ± SEM.

Following seizure elicitation, both groups exhibited a prolonged period of hypoxia which had comparable levels of overall hypoxic burden defined by the area below 10mmHg (Figure 1C). Notably, Cx40^-^/^-^ mice displayed an abnormal neurovascular response in the immediate postictal period at 3–4 minutes (Figure 1E; independent-samples t-test, p < 0.05). Both groups showed a characteristic immediate postictal pO_2_ profile with two distinct rebounds. Compared with C57BL/6J mice, Cx40^-^/^-^ mice exhibited a significantly longer delay to the second peak following stimulation (Figure 1F; independent-samples t-test, p < 0.01). However, hippocampal pO_2_ at the second peak was similar between C57BL/6J controls and Cx40^-^/^-^ mice (Figure 1G). Collectively, these data suggest that Cx40 influences the dynamics of postictal oxygen recovery, prompting further consideration of the role of vascular gap junctional signaling in seizure-induced cerebrovascular dysfunction.

## Discussion

Genetic deletion of Cx40 altered the temporal profile of postictal severe hypoxia without affecting seizure duration. Brief focal hippocampal seizures produced prolonged reductions in local pO_2_ in both genotypes, consistent with previous studies showing that electrically evoked seizures induce severe postictal hypoxia, with tissue pO_2_ values falling below 10 mmHg for more than an hour after seizure termination^6,14^. Furthermore, because postictal hypoxia severity directly correlates with seizure duration^6^, the absence of genotype-dependent differences in seizure duration (Figure 1A) indicates that the altered hypoxic profile in Cx40 knockout mice is more likely attributable to vascular rather than seizure-related mechanisms.

A notable finding was that Cx40^-^/^-^ mice experienced greater hypoxia immediately after seizures and a delayed second phase of oxygen recovery, despite reaching similar peak postictal pO_2_ levels as wild-type animals (Figure 1C-G). These data indicate that loss of Cx40 alters the temporal profile of postictal oxygen dynamics without preventing eventual recovery of tissue oxygenation. Previous work has shown that postictal hypoperfusion/hypoxia is driven by sustained vasoconstriction mediated through neuronal COX-2 signaling and downstream vascular calcium-dependent pathways^6,10^. Our findings are therefore consistent with the possibility that endothelial Cx40-dependent endothelial signaling influences the efficiency of vascular recovery during the immediate postictal period. This interpretation is supported by the established role of Cx40 in vascular signaling. Cx40 is exclusively expressed in the endothelial cells of mice^12^ and contributes to vascular homeostasis, blood-pressure regulation, and the conduction of vasomotor responses along arterial networks^11,12,15^. Disruption of endothelial gap-junction signaling through genetic deletion of Cx40 is known to impair conducted vascular responses and to compromise blood-flow homeostasis after experimental stroke in mice^12^. Consequently, loss of Cx40 could plausibly alter physiological mechanism governing postictal oxygen recovery; however, because blood flow was not directly measured, the relative contributions of vascular and metabolic factors remain to be determined.

Although behavioral outcomes and blood-brain barrier integrity were not examined directly, altered postictal hypoxia dynamics may have important downstream consequences. Postictal hypoperfusion/hypoxia contributes to transient cognitive and motor impairment^6^, while postictal hypoperfusion has also been identified in brainstem respiratory centers implicated in SUDEP risk^16^. Furthermore, disruption of vascular integrity and neurovascular coupling is increasingly recognized as a contributor to epilepsy associated pathology^17–19^. In this context, the present study provides initial evidence that Cx40-dependent vascular signaling influences the temporal oxygen dynamics of postictal hypoxia. Although the use of a global Cx40 knockout model, a single focal hippocampal seizure paradigm, and tissue oxygen measurements rather than direct assessments of vascular diameter or blood flow limit mechanistic interpretation, genetic deletion of Cx40 did not alter seizure duration and was associated with greater early hypoxia and delayed recovery, supporting the concept that postictal neurovascular responses can be regulated independently of seizure activity itself. While Cx40 is generally regarded as an endothelial connexin, studies have identified Cx40 expression in vascular smooth muscle cells in specific vascular beds and experimental conditions^20–22^, suggesting that non-endothelial mechanisms^23^ cannot be excluded in the present study. These findings raise the possibility that Cx40 dependent signaling within the cerebrovascular network contributes to the regulation of postictal oxygen dynamics following seizures and that its disruption may exacerbate neuronal stress and vascular dysfunction within seizure-prone tissue. Future studies using cell type-specific connexin manipulations, chronic epilepsy models, and direct assessments of cerebrovascular dynamics will be required to determine whether preservation of Cx40 dependent signaling can mitigate the long-term consequences of recurrent postictal hypoperfusion/hypoxia.

## Funding and Author Contributions

This work was supported by an operating grant from CHIR (159667) to D.G.W, an operating grant from CHIR (130495) to G.C.T., postdoctoral fellowships from University of Calgary (Eyes High), Alberta Innovates Health Solutions (AIHS), Canadian Institutes of Health Research (CIHR), a startup grant from the Faculty of Medicine, Memorial University and a New Investigator Award from Heart and Stroke Foundation of Canada (NI-24-0036510) to A.Z.

D.G.W. and A.Z. conceived the study. J.S.F. performed the experiments and conducted the data analysis. D.G.W., J.S.F., G.C.T., and A.Z. contributed to the writing and revision of the manuscript. A.Z. finalized the manuscript. All authors reviewed and approved the final version of the manuscript.

